# Bacterial colonisation dynamics of household plastics in a coastal environment

**DOI:** 10.1101/2021.11.02.466905

**Authors:** Luke Lear, Daniel Padfield, Tirion Dowsett, Maia Jones, Suzanne Kay, Alex Hayward, Michiel Vos

## Abstract

Accumulation of plastics in the marine environment has widespread detrimental consequences for ecosystems and wildlife. Marine plastics are rapidly colonised by a wide diversity of bacteria, including human pathogens, posing potential risks to human health. Here, we investigate the effect of polymer type, residence time and estuarine location on bacterial colonisation of common household plastics, including pathogenic bacteria. To do so, we submerged five main household plastic types: low-density PE (LDPE), high-density PE (HDPE), polypropylene (PP), polyvinyl chloride (PVC) and polyethylene terephthalate (PET) at an estuarine site in Cornwall (U.K.) and tracked bacterial colonisation dynamics. Using both culture-dependent and culture-independent approaches, we found that bacteria rapidly colonised plastics irrespective of polymer type. While biofilm community composition changed with colonisation time, no difference was observed between polymer types. Likewise, the presence of pathogenic bacteria, quantified using the insect model *Galleria mellonella*, increased over a five-week period, with no consistent differences observed between polymer types. Pathogens isolated from plastic biofilms using *Galleria* enrichment included *Serratia* and *Enterococcus* species and harboured a wide range of antimicrobial resistance genes. Our findings show that plastics in coastal waters are rapidly colonised by a wide diversity of bacteria, including known human pathogens, independent of polymer type.

## Introduction

Plastics pollution has huge negative impacts on marine environments (1). In 2017, around 348 million tonnes of plastic were produced globally (2). This figure is increasing year on year (2), with a significant percentage of all plastics produced ultimately finding their way into the marine environment (3, 4). Plastics have multiple adverse effects on a wide diversity of wildlife, including entanglement, ingestion and physical damage (e.g. (5-8)). Moreover, it is increasingly recognised that there is potential for plastic pollution to pose a threat to human health (9), from the effects of ingestion of micro- and nano particles (e.g. via shellfish or drinking water (10)) followed by cellular uptake (11), to decreased psychological wellbeing through pollution of the natural environment (12).

The surface of plastic fragments in aquatic environments is rapidly colonised by microorganisms, leading to diverse biofilm communities distinct from that of the surrounding seawater (13-15). Bacterial colonisation of marine plastics is dependent on a variety of abiotic factors, including season, location, colonisation time and polymer type (13, 16, 17). A growing number of studies have used 16S amplicon sequencing approaches to specifically investigate bacterial colonisation of different substrates. Clear differences in bacterial community composition between plastic and other surfaces (e.g. wood or glass) have been reported (e.g. (18, 19)) (but see (20)). Differences in plastic colonisation based on beta-diversity metrics have been mixed. Differences have been reported between polystyrene (PS) and polyethylene (PE) and polypropylene (PP) (21, 22), between PS and PP, PE and polyethylene terephthalate (PET) (23) and between PE and PET (24).However, other studies have failed to find biofilm community differences between plastics, e.g. between polyethylene (PE), polypropylene (PP) and polystyrene (PS) (18), between PE and PS (20, 25) or between PE and PP (19).

Among bacterial colonisers of marine plastics are pathogenic species known to cause disease in animals (e.g. (26, 27)) and humans (e.g. (28-30)). Plastics could serve as potential vectors for pathogens (30) and/or bacteria carrying antimicrobial resistance genes (31). Plastic pollution could also result in elevated horizontal gene transfer by facilitating close contact (32), and plastic-sorbed contaminants (33) could potentially speed up antimicrobial resistance development (34). Therefore, in addition to direct threats related to ingestion, plastic-borne bacteria may also pose an infection risk to human and wildlife health (35-37). However, previous work has concentrated on indirect quantification of potential pathogens using agar cultivation or amplicon sequencing which cannot quantify the virulence or pathogenicity of bacteria,

The aim of this study is to investigate the colonisation dynamics of bacterial pathogens on five of the most common types of household plastics (low-density PE (LDPE), high-density PE (HDPE)), polypropylene (PP), polyvinyl chloride (PVC) and polyethylene terephthalate (PET)), at multiple coastal sites, and over time. We focus on plastic colonisation in coastal waters where terrestrial inputs of plastics (17) and human-, animal- and soil-associated pathogens is highest (38). We investigate bacterial colonisation dynamics on plastics incubated in the marine environment by quantifying density for subsets of cultivable diversity and testing for differences in taxonomic diversity via 16S amplicon sequencing. We directly tested for the presence of pathogens on biofilm plastics, employing a recently developed selective isolation method using the *Galleria mellonella* insect virulence model (39). We performed two separate experiments: (i) we measured bacterial colonisation after seven weeks across three different locations, quantifying bacterial colonisation through cultivation-based approaches and measuring the presence of pathogens using the *Galleria* assay; (ii) we focused on one location close to shore and in addition to cultivation and the *Galleria* assay, used 16S rRNA amplicon sequencing to quantify bacterial colonisation over time.

## Materials and Methods

### Plastics

Five plastic types were collected from household recycling: low-density polyethylene (LDPE) from sandwich bags, high-density polyethylene (HDPE) from milk bottles, polypropylene (PP) from peanut butter tubs, polyvinyl chloride (PVC) from roofing material and polyethylene terephthalate (PET) from drink bottles. Plastics were cut into 85mm by 25mm strips with a 5mm diameter hole made at one end using a holepunch. One sample of each type of plastic was attached to a large cable tie via smaller, colour coded cable ties to form a ring (Figure 1a). Rings were then attached to metal mesh bags using cable ties (Figure 1b). Plastics were sterilised using 70% ethanol before environmental incubation.

**Figure 1.**
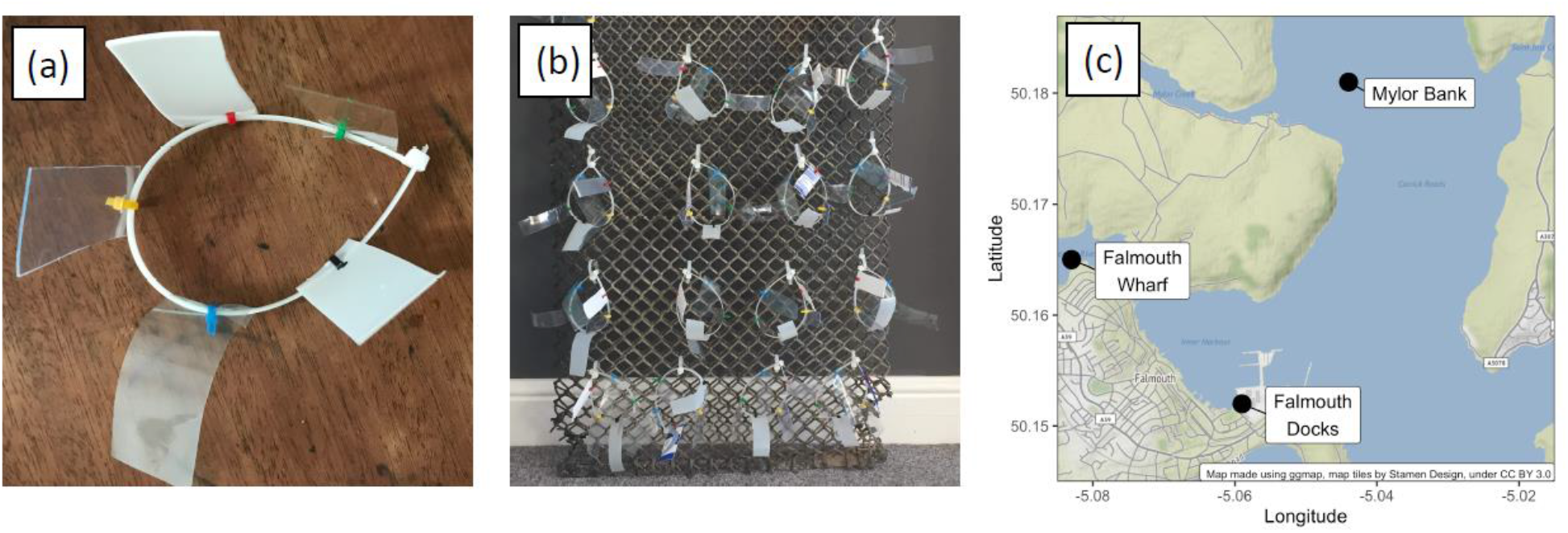
Overview of the methods used to submerge household plastics in the Fal Estuary. (a) Strips of LDPE, HDPE, PP, PET and PVC strips were attached to a big cable tie ring using smaller, colour coded cable ties. (b) Each large ring was attached to a metal mesh bag using cable ties which was then submerged at each site (c) Sites used in this study: Falmouth Docks, Falmouth Wharves and Mylor Bank. All three sites were used in the first ‘Location’ experiment whereas the second ‘Colonisation’ experiment occurred only at the Falmouth Docks site.

### Location experiment

A total of 18 rings (attached to three mesh bags) were deployed on 10th December 2019 at three sampling sites in the Fal Estuary (Falmouth, Cornwall, UK): Falmouth Docks (50.152, -5.059), Falmouth Wharves (50.165, -5.083) and Mylor Bank (50.181, -5.044) (Figure 1c) with help from Cornwall Port Health Authority. Mylor Bank was further from the shore and was further away human habitation (Figure 1c). The bags were attached to pontoons at Falmouth Wharves and Falmouth Docks and to a buoy at Mylor Bank using a rope, ensuring the bags were fully submerged. Plastic rings were retrieved from each site on 28th January 2020 (after 7 weeks) and placed separately in a sterile plastic box for transportation. Sample processing commenced within one hour of collection.

### Colonisation experiment

To measure differences in the bacterial colonisation of different plastic types through time, 28 rings of plastics (four timepoints, seven replicates) were attached to a single metal mesh bag and submerged as described above at the Falmouth Docks site on February 4^th^, 2020. One set of plastic rings were collected on February 11th, 18^th^ and 24^th^ and March 3^rd^ (weeks 1, 2, 3 and 5) and brought back to the lab.

### Biofilm processing

Under sterile conditions, each plastic strip was cut in half using sterilised scissors and placed in a 50mL falcon tube containing 10mL sterile NaCl buffer (9g/L; Fisher Chemicals, Loughborough, UK) and five sterile glass beads to facilitate removal of biofilm (Millipore Colirollers Plating Beads, Billerica MA, USA). Each tube was vortexed for 60 seconds at 2500 r.p.m. to remove and suspend plastic biofilms. As our focus was on bacteria of terrestrial origin, specifically potential human pathogens, serial dilutions of biofilm suspensions were plated on LB agar (Fisher BioReagents, Loughborough, UK) and coliform agar (Millipore Sigma, Billerica MA, USA) and incubated at 37°C. LB agar selects for relatively fast-growing bacteria; coliform agar was used to quantify both coliform bacteria and *E. coli*. Colony forming units (CFU) were counted after 24 hours of incubation at 37°C. All counts were standardised to CFU/cm^2^. Biofilm suspensions were stored in glycerol (Fisher Chemicals, Loughborough, UK) (20% final concentration) at -80°C.

### *Galleria mellonella* virulence assay

*G. mellonella* larvae were purchased from Livefood U.K. (http://www.livefood.co.uk) and used within one week of purchase. A 100μL Hamilton syringe (Sigma-Aldrich Ltd, Gillingham, UK) with 0.6 × 30mm needles (BD Microlance 3, Becton Dickinson, Plymouth, UK) was used to inject the larvae with 10μL of defrosted biofilm freezer stock into the last left proleg, using 20 larvae (location experiment) or 10 larvae (colonisation experiment) for each sample. The larvae were anaesthetised on ice for 30 minutes before injection. Needles were sterilised between samples by flushing with 70% ethanol followed by NaCl buffer. Two negative controls for the experiment were used: a buffer control using 10μL of sterile NaCl to control for the impact of physical trauma, and a no-injection control to account for background larvae mortality. After injection, larvae were incubated at 37°C and inspected at 24-, 48- and 72-hours post-injection (location experiment) or 24- and 48 hours post-injection (colonisation experiment) to record mortality. Larvae were scored as dead if they did not respond to touch stimuli (39).

### Isolation of pathogenic bacteria

To isolate pathogens causing *G. mellonella* mortality, we reinjected 20 of the most virulent communities into ten larvae as described above. Larvae demonstrating melanisation – a key indicator of infection – were anesthetised by placing on ice before their haemocoel was extracted. To extract haemocoel, 70% ethanol was used to sterilise the area around the last left proleg before the proleg was removed using sterile micro-scissors and the haemocoel collected using a pipette. This method is advantageous over whole larvae destruction as it minimises contamination with skin and gut bacteria (Hernandez et al 2019). Collected haemocoel (∼5-15μL) was diluted in 500μL of buffer, plated onto LB agar and incubated for 24 hours. Colonies were then picked and grown in 750μL of LB at 37^º^C for 24 hours. Cultures were frozen at -80^º^C in glycerol (at a final concentration of 25%). Defrosted stocks were diluted to (1 × 10^5^ CFU/mL) and injected into ten larvae which were then incubated for 18 hours; deceased larvae confirmed that isolated clones were pathogenic and not commensal gut or skin bacteria.

### 16S rRNA amplicon sequencing

DNA was extracted from biofilms using a DNeasy UltraClean Microbial Kit (Qiagen, Hilden, Germany) according to the standard protocol with the only modification to increase the initial centrifugation speed and time (10K x g for 10 minutes) to pellet bacteria suspended in the 20% glycerol stock. DNA concentrations were quantified using the Qubit HS DNA kit (Invitrogen), purity was assessed using nanodrop 260:280 ratios, and integrity was assessed using a 1% agarose gel. A 251bp conserved fragment in the V4 hypervariable region was targeted using N515f and N806r primers (https://earthmicrobiome.org/protocols-and-standards/16s/) with phasing and a pool of indexed primers suitable for multiplex sequencing with Illumina technology. Sequencing was performed using an Illumina MiSeq V2 500 by the University of Exeter Sequencing Service. Sequencing adapters and any bases below a score of Q22 were removed, alongside any reads <150 bp using ‘*Cutadapt*’ (40). We then processed and analysed the sequence data in R (v4.0.3) (41) using the packages ‘*dada2*’ (42) and ‘*phyloseq*’ (43). Following the standard full-stack workflow, we estimated error rates, inferred and merged sequences, constructed a sequence table, removed chimeric sequences and assigned taxonomy. During processing, the first 25bp of forward and reverse reads were trimmed. Taxonomies were assigned to amplicon sequence variants (ASVs) using the SILVA database (44). We estimated phylogeny using ‘*fasttree*’ (45) to allow for the calculation of Unifrac distances (which take into account the phylogenetic distance between ASVs) between communities. We then removed any reads that had not been assigned to at least the phylum level (613 of 12564 unique ASVs) and any sequences assigned to the phylum Cyanobacteria. Processing and filtering steps resulted in three samples containing fewer than 10,000 reads being removed (one replicate from week 1 LDPE, one replicate from week 1 HDPE, and one replicate from week 2 of PP), with the remaining samples having a maximum read number of 673,525, a minimum of 10,364 and a median of 24,920.

### Whole Genome Sequencing

DNA isolation, Illumina HiSeq sequencing and basic bioinformatics were performed through MicrobesNG, Birmingham, UK. Vials containing beads inoculated with liquid culture were washed with extraction buffer containing lysostaphin and RNase A, and incubated for 25 minutes at 37°C. Proteinase K and RNaseA were added and vials were incubated for a further 5 minutes at 65°C. Genomic DNA was purified using an equal volume of SPRI beads and resuspended in EB buffer. DNA was quantified in triplicate using the Quantit dsDNA HS assay in an Eppendorff AF2200 plate reader. Genomic DNA libraries were prepared using the Nextera XT Library Prep Kit (Illumina, San Diego, CA, USA) following the manufacturer’s protocol with the following modifications: two nanograms of DNA instead of one were used as input, and PCR elongation time was increased to 1 minute from 30 seconds. DNA quantification and library preparation were carried out on a Hamilton Microlab STAR automated liquid handling system. Pooled libraries were quantified using the Kapa Biosystems Library Quantification Kit for Illumina on a Roche LightCycler 96 qPCR machine. Libraries were sequenced on the Illumina HiSeq using a 250bp paired end protocol.

Sequencing reads were adapter trimmed using ‘*Trimmomatic 0*.*39*’ with a sliding window quality cutoff of Q15 (46). De novo assembly was performed on samples using ‘*Unicycler v0*.*4*.*8*’ (47), and summary stats of assemblies were calculated using CheckM v1.1.3 (48). Genome assemblies were visualised using Bandage (49) and split when there were two large non-contiguous assemblies which were then run separately through the other stages. The assemblies which were likely two or more species were identifiable by a high contamination value (∼100%). Contigs were annotated using Prokka 1.14.6 (50). Taxonomic classification of our assemblies was performed using ‘*MMSeqs2 13*.*45111*’ (51) using the GTDB database (52), and of 16S gene amplicons using ‘*dada2’* (53) using the SILVA reference database (44). Assemblies were screened for antimicrobial resistance genes and virulence factors using ‘*AMRFinderPlus v3*.*10*.*1*’ (54).

### Statistical analyses

#### Analysing differences in culturable abundance between plastics, sites, and time

All analyses were conducted using the statistical programming language R v4.0.3 (41) and all graphs were produced using the ‘*ggplot2*’ package (55). Linear models were used to examine differences in culturable bacterial abundance between locations. The response variable was log_10_ abundance+1 cm^-2^ plastic to normalise the residuals, and the predictors were location, plastic type and their interaction. Separate models were run for each category of culturable bacteria (bacteria culturable on LB, coliform and *E. coli*). Model selection was performed using using likelihood ratio tests. Pairwise multiple comparisons to examine significant differences among locations or plastics were performed using the R package ‘*emmeans*’ (56). Changes in abundance through time during colonisation were analysed using linear models. The response variable was again log_10_ abundance+1 cm^-2^ plastic with week of sampling as a continuous predictor and type of plastic as an interaction term. Separate linear models were run for all three categories of culturable bacteria. Model selection and post-hoc multiple comparisons were performed as above.

### Analysing virulence of plastisphere biofilms

Survival of *Galleria* inoculated with bacteria isolated from biofilms was quantified as a measure of virulence of the plastisphere communities. Survival curves were fitted using Bayesian regression in the package ‘*rstanarm*’ (57) and parameters were estimated using ‘*tidybayes*’ (58). Compared to the popular Cox model, Bayesian implementation can more easily visualise uncertainty and better handle random effects. For survival curves of *Galleria* injected with bacteria isolated from 7-week-long biofilms, we fitted a proportional hazards model with an M-splines baseline hazard, with site, plastic, and their interaction as fixed effects and a random effect of biofilm, as multiple *Galleria* were all inoculated with the same biofilm. For the survival curves through time at Falmouth Docks, we fitted the same model, but with week, plastic, and their interaction as fixed effects and a random effect of biofilm. Models were run for 3000 iterations and three chains were used with uninformative priors. Model convergence was assessed using Rhat values (all values were 1) and manual checking of chain mixing. For both models, log hazards were estimated for each plastic in each site or week. To examine differences in virulence among sites and plastics, hazard ratios were calculated as the exponential of the difference between two log hazards. A hazard ratio above one indicates an increase in virulence compared to the baseline treatment, with a value below one indicating a decrease in virulence compared to the baseline. Median log hazard across treatments was used as the baseline. We calculated median hazard ratios with 95% credible intervals and the probability that the given hazard ratio was above 1. Median hazard ratios with 95% credible intervals that do not cross 1 indicate a significant difference in virulence between two factors. For visualisation purposes, we show the hazard ratio of each location/polymer type where the baseline is the median log hazard in each experiment, allowing differences in virulence between sites/polymer types to be easily seen.

### Analysing plastisphere community composition between different plastic types

To investigate how plastisphere bacterial communities changed, we examined changes in community composition, alpha and beta diversity through time. We used weighted Unifrac distance (59) as a measure of compositional dissimilarity between communities, which weights branches in a phylogeny based on the relative abundance of each ASV. As this distance requires a rooted phylogeny, we rooted the tree based on the longest tree branch terminating in a tip. Differences in composition between plastic communities were analysed using the R packages ‘*phyloseq*’ (43) and ‘*vegan*’ (60). We tested whether week of colonisation and plastic type altered community composition using permutational ANOVAs. Permutational ANOVA tests were run using ‘*vegan::adonis*’ with 9999 permutations and differences in group dispersion (which test for differences in beta diversity between treatments) were analysed using ‘*vegan::betadisper*’. An overall model including plastic type and colonisation week as interacting factors was performed first. We then split the data to examine an overall effect of plastic type by performing a separate PERMANOVA for each week and performing pairwise PERMANOVAs between each plastic type to assess differences. P values were corrected using the *fdr* method (61). When looking at differences in beta diversity, we calculated the distance from the centroid of each plastic by week combination, and then ran a linear model where distance from the centroid was the response variable, and plastic and week were potentially interacting factors. Model selection was performed as above.

To investigate differences in alpha diversity and evenness between communities, we rarefied data so that all samples had the same number of reads (11,045). Richness (alpha diversity) was taken as the total number of ASVs and evenness was calculated as Pielou’s evenness (62). The amount of change and variation in the number of ASVs across weeks and across plastic types was tested using linear models. Total ASVs were log_10_-transformed to normalise residuals, with plastic type and week as potentially interacting predictors. As we did not expect alpha diversity to necessarily change linearly across weeks, we included it as a categorical predictor. For evenness, the analysis was the same, but without any transformation on the response. Model selection was performed as stated earlier.

## Results

### Bacteria abundance was lowest on plastics further away from human habitation

To determine the effects of location and polymer type on bacterial colonisation, five different plastics were incubated at one-meter depth at three estuarine locations near Falmouth, U.K. After seven weeks of submergence, the abundance of bacteria culturable on LB, coliforms, and *E. coli*, significantly differed between locations (ANOVA between models with and without location as a predictor: bacteria culturable on LB: F_2,83_ = 69.32, p < 0.001; coliforms: F_2,83_ = 54.34, p < 0.001; *E. coli*: F_2,83_ = 14.44, p < 0.001) (Figure 2). Across the three quantified groups, abundance was consistently highest at Falmouth Dockyard, which is located closest to human habitation (bacteria culturable on LB: mean =724 CFU cm^2^, 95%CI = 490 - 1096; coliform: mean =50 CFU cm^2^, 95%CI = 30 – 83 *E. coli*: mean =3.3 CFU cm^2^, 95%CI = 2.4 – 4.6) and lowest at Mylor Bank which is furthest from the shore and human habitation (bacteria culturable on LB: mean =25 CFU cm^2^, 95%CI = 17 – 38; coliform: mean =1.2 CFU cm^2^, 95%CI = 0.7 – 2.1; *E. coli*: mean =1.04 CFU cm^2^, 95%CI = 0.76 – 1.4), although not all contrasts were significant (Table S1).

**Figure 2.**
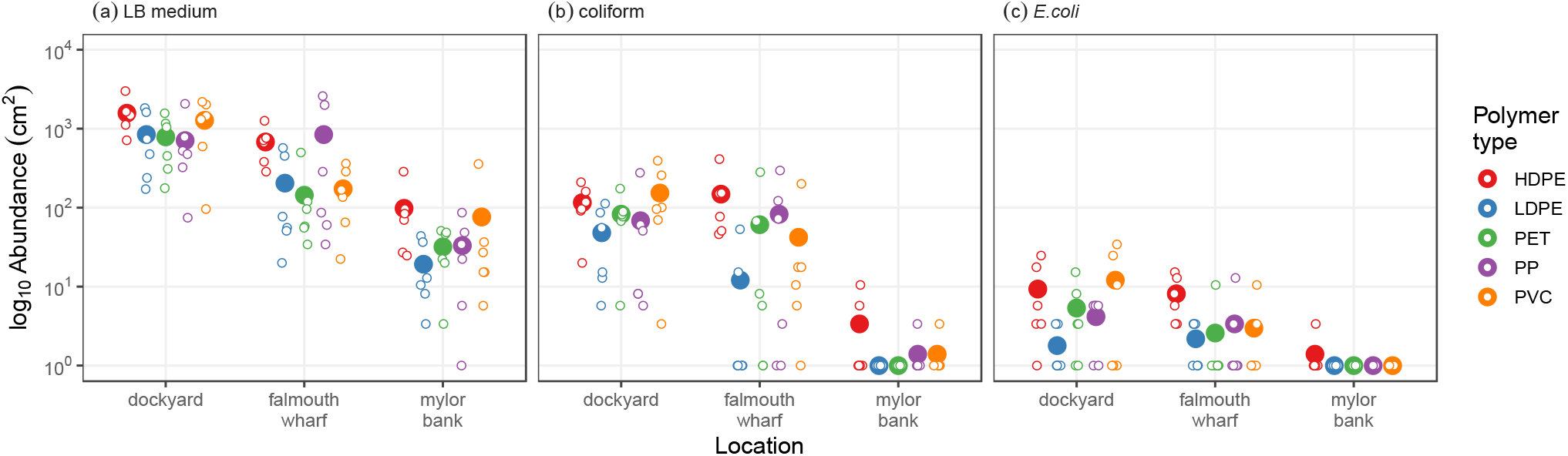
Abundance of (a) bacteria on LB medium, (b) coliform bacteria and (c) *E. coli* found across polymer types and locations. Large points represent mean abundances of plastics at each site and small points represent individual replicates.

Significant differences in abundance were found between plastics (ANOVA between models with and without plastic as a predictor: bacteria culturable on LB: F_4,83_ = 4.65, p = 0.002; coliforms: F_4,83_ = 4.05, p = 0.005; *E. coli*: F_4,83_ = 3.11, p = 0.019), but there was no significant interaction between location and plastic (ANOVA between models with and without interaction between plastic and location: p > 0.05 for LB, coliform and *E. coli*). Abundance was consistently highest on HDPE, but this was only significant across all three quantified groups when compared to LDPE (Tukey comparison between HDPE and LDPE: bacteria culturable on LB: t = 3.86, p = 0.002; coliform: t = 3.88, p = 0.0018; *E. coli*: t = 3.31, p = 0.0118; Table S1).

### Rate of bacterial colonisation did not differ between polymer types

To investigate bacterial colonisation of plastics through time, we performed a second experiment at the Falmouth Docks site only, and sampled plastics after one, two, three and five weeks of submergence. During this period, the abundance of bacteria growing on LB, coliforms, and *E. coli* all increased (slope of increase in abundance in best model: bacteria culturable on LB: 0.23, 95%CI = 0.17-0.29; coliform = 0.32, 95%CI = 0.25-0.39; *E. coli*: 0.17, 95%CI = 0.13-0.19) (Figure 3). For both bacteria culturable on LBand coliforms, the increase in abundance occurred consistently across all types of plastic (ANOVA with and without plastic as a predictor: all p values > 0.05). The only difference in colonisation rate between plastics was found for *E. coli* (ANOVA with and without the interaction between week and plastic: F_4,130_ = 5.95, p = 0.0002), where the rate of colonisation was lowest on LDPE (slope of increase in abundance in best model: *E. coli* on LDPE: 0.05, 95%CI = -0.02 – 0.12), but this was only significantly lower than PET (Tukey comparison between slope of LDPE and PET: t = -4.12, p = 0.0005) and PVC (Tukey comparison between slope of LDPE and PVC: t = -3.92, p = 0.013).

**Figure 3.**
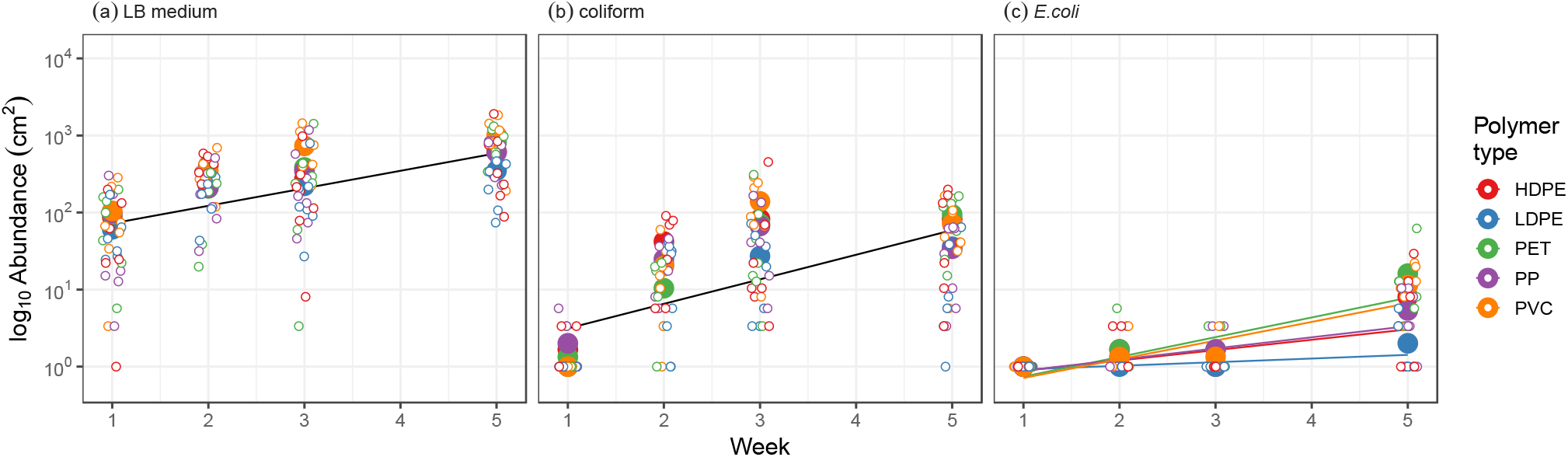
Abundance of (a) bacteria on LB medium, (b) coliform bacteria and (c) *E. coli*, through five weeks of colonisation at the Falmouth Dockyard site. Small points represent individual replicates and large points represent mean abundances on each plastic at each week. Lines represent the predicted best fit from the model (see Methods), with the single black line in (a) and (b) indicating that there is no effect of plastic on abundance.

### Plastic biofilm community composition changes through time, but does not differ between polymer types

In addition to culture-dependent approaches, we performed 16S amplicon sequencing across all four timepoints and plastic types for the colonisation experiment. Alpha diversity increased during the first two weeks of colonisation, after which the number of unique ASVs remained stable between weeks 2 and 5 (Figure 4a). This same pattern was seen in the evenness of the biofilm communities, with abundances being more even across species in week 1 compared to weeks 2 to 5 (Figure 4b) which is consistent with a scenario where a subset of taxa increases in frequency over the course of plastic colonisation.

**Figure 4.**
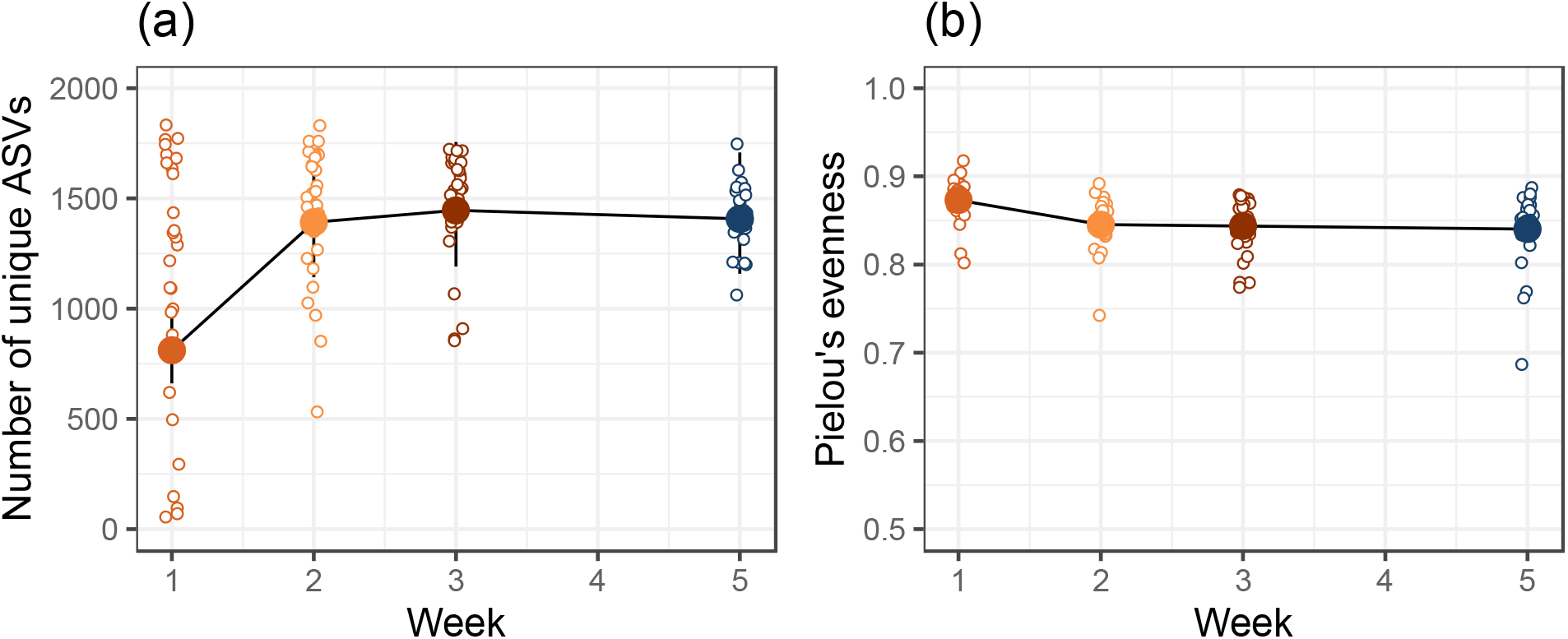
Change in (a) alpha diversity and (b) evenness of plastic biofilm communities. (a) Alpha diversity increased from week 1 to week 2 and then remained stable. (b) At the same time, communities became more even, with the only difference in evenness occurring between week 1 and 2. In both panels, small points are individual replicates, large points and line bars represent estimates and 95% confidence intervals from the best model respectively (see Methods), and model estimates are joined by lines to help visualise differences between weeks.

Bacterial community composition changed significantly over the course of the experiment (PERMANOVA, F_3,108_ = 23.84, R^2^ = 0.38, p = 0.001) (Figure 5). The first principal coordinate explained 22.0% of the total variation and separated the weeks, with the greatest difference between week 1 and week 5. While there was no interaction between plastic and week, plastic type significantly altered community composition when added alongside week (PERMANOVA, F_4,108_ = 1.85, R^2^ = 0.04, p = 0.013), although the amount of variation it accounted for was very small (∼4%). To further examine this effect, we ran multiple pairwise permutational ANOVAs on the entire dataset and within each week. We found no significant differences between any two plastics on the whole dataset or within each week (PERMANOVAs: all p_adj_ > 0.05), indicating no consistent effects of plastic type. There were no overall differences in between-community diversity (beta-diversity) across weeks or plastics, with the simplest model being one with no predictor (Table S2).

**Figure 5.**
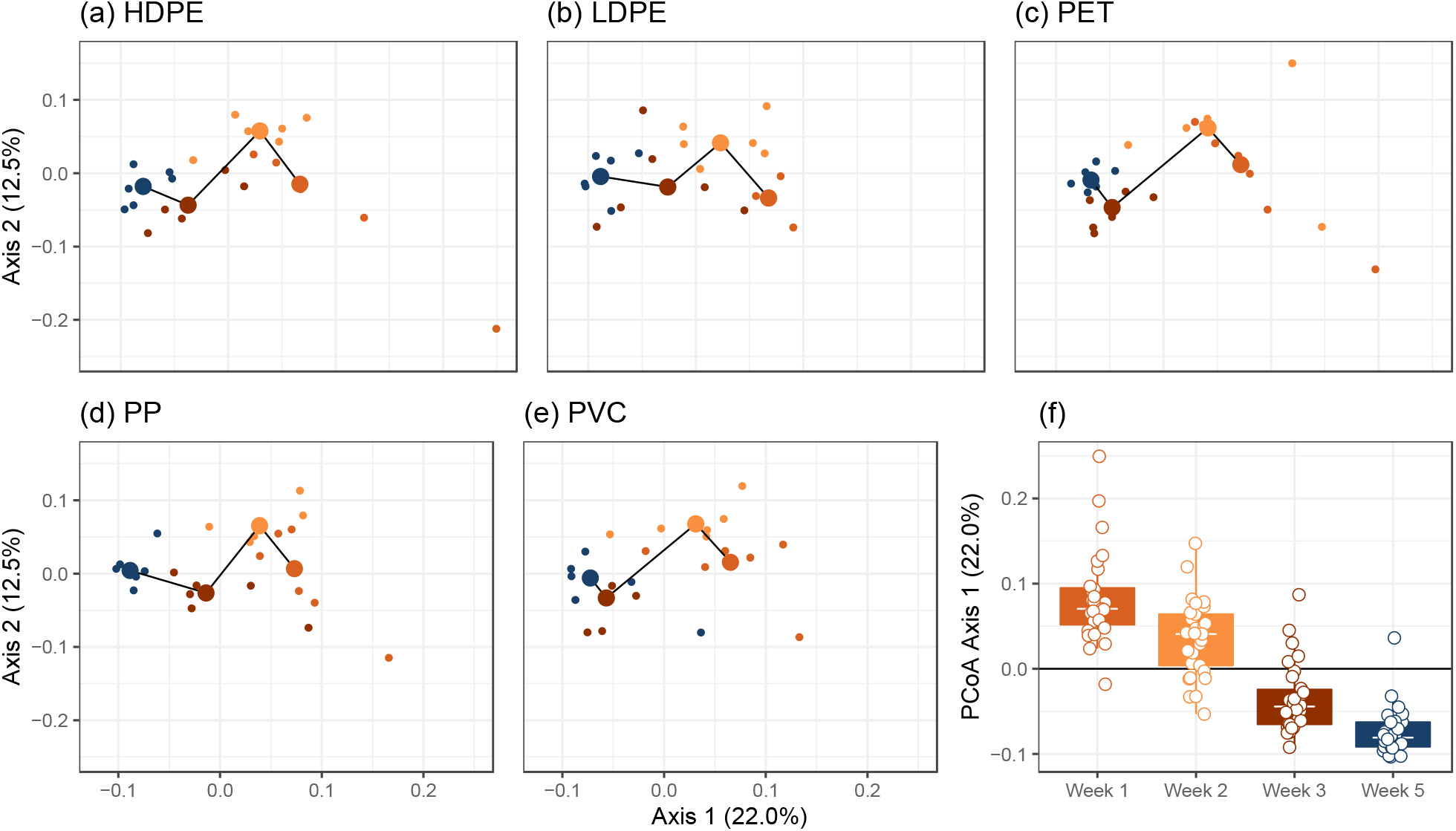
Changes in community composition through time (different colours) and between plastics (panels a-e). (a-e) Principal Coordinate (PCoA) plot of communities based on weighted-Unifrac distance. Each panel represents biofilms from different polymer types and shows that community composition changes consistently through time irrespective of plastic type. The percentage of variation explained is shown on each axis (calculated from the relevant eigenvalues). (f) The separation of communities through time along PCoA axis 1 scores. In (a-e) small points are individual communities and large points are positions of centroids, with the black lines showing how centroid position changes through time. In (f) points are individual communities, tops and bottoms of the bars represent the 75^th^ and 25^th^ percentiles of the data. White lines are the medians, and the whiskers extend from the respective hinge to the smallest of largest value no further than 1.5*interquartile range.

### Virulence of plastic biofilm communities

We employed a recently developed method where pathogenic bacteria from microbiome samples are selectively enriched using the *G. mellonella* wax moth larva virulence model (39). In the location experiment, all hazard ratios between the different sites had credible intervals that crossed one, indicating non-significance at a traditional 0.95 level. However, biofilms from Falmouth Dockyard (mean time to death = 28.5 hours; proportion that died = 0.48) and Falmouth Wharf (mean time to death = 30.8 hours; proportion that died = 0.48) were 2.5 times more virulent than biofilms from Mylor Bank (mean time to death = 34.4 hours; proportion that died = 0.32) (Figure 6a, Table S3). There was an 87% probability that the Dockyard and Wharf communities were more virulent than those from Mylor bank for both locations (Table S3).

**Figure 6.**
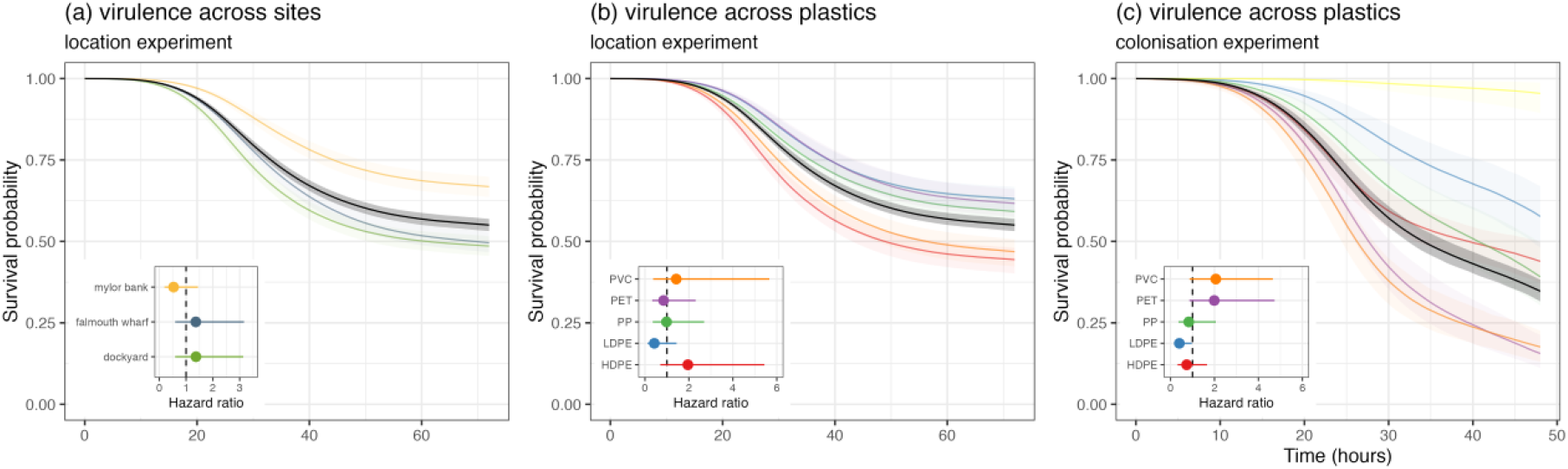
Survival curves of *G. mellonella* inoculated with plastic biofilm communities from (a, b) the location experiment and (c) the colonisation experiment. Across all panels, the black line and shaded region depicts the median virulence of a biofilm community from that experiment, and the median log hazard was used as the baseline for calculating hazard ratios. (a) Effect of location on virulence of biofilm communities. (b) Effect of polymer type on virulence of biofilm communities from the location experiment. (c) Effect of polymer type on virulence of biofilm communities from the colonisation experiment. Lines represent the median prediction and shaded bands represent 95% credible intervals of those predictions. Inset plots are the hazard ratio where the median hazard of the experiment is the baseline. In (c) bright yellow is the survival curve for *G. mellonella* injected with water.

In the colonisation experiment, biofilms from weeks 1, 2, and 3 killed only 4, 7, and 11% of *Galleria* respectively. In contrast, biofilms from week 5 were much more virulent, with 65% of *Galleria* dying during the assay. In both the location and colonisation experiments, we looked for differences in virulence between polymer types (Figure 6a). To classify a polymer type as having more or less virulent communities than others, it would need to have a higher (or lower) hazard ratio than average in both experiments (Table S3). There were no hazard ratios between polymer types that had 95% credible intervals that did not overlap with 1 for both experiments meaning that there were no significant differences in virulence between polymer types. However, we found evidence that LDPE communities are less virulent than other polymer types, with all hazard ratios being above 1 when LDPE was the baseline polymer type (Table S3), with an 89% probability (on average) of the other polymer type being more virulent. The four other polymer types did not have consistent differences in their hazard ratios.

### The identity of putative human pathogens

To obtain a more detailed picture of the taxonomic identity and antimicrobial resistance of individual pathogens in our plastic biofilm samples, we genome-sequenced 16 bacterial clones isolated from some of the most virulent biofilm communities spanning different locations and plastic types (Table 1). Eleven genome assemblies had high completion and low contamination and could be unambiguously assigned to a single bacterial species, but this was not the case for five remaining genome samples – which must have been due to the co-isolation of multiple clones. To retain as much information as possible, these assemblies were not discarded but were bioinformatically separated using the genome viewer Bandage, with assemblies split into non-contiguous sections (46). Using this approach, four contaminated genomes could be split into eight genomes (2 & 3, 6 & 7, 8 & 9, 15 & 16; Table 1). One remaining assembly (17; Table 1) could not be split because it only had one contiguous genome. Retaining a large chimeric assembly was also the case for (split) sample 16 and so for both assemblies multiple identities are presented instead of one (Table 1).

**Table 1.** Genome characteristics of 20 clones isolated from plastics using the *Galleria* enrichment method.

| isolate | plastic | site | contig taxonomic assignment | 16S taxonomic assignment | completeness | contamination | GC | genome size (bp) | number of contigs | N50 | number of AMR genes |
| --- | --- | --- | --- | --- | --- | --- | --- | --- | --- | --- | --- |
| 1 | HDPE | falmouth wharf | *Serratia liquefaciens* | *Serratia* | 100.00 | 0.89 | 0.55 | 5,549,824 | 78 | 327,206 | 2 |
| 2 | HDPE | falmouth wharf | *Serratia* | *Raoultella/Enterobacter/Serratia* | 99.86 | 0.15 | 0.59 | 5,222,268 | 41 | 2,939,895 | 6 |
| 3 | HDPE | falmouth wharf | *Bacillus licheniformis* | *Bacillus licheniformis* | 98.82 | 0.00 | 0.46 | 4,254,100 | 20 | 2,234,065 | 1 |
| 4 | HDPE | mylor bank | *Serratia* | *Serratia marcescens* | 99.86 | 0.15 | 0.59 | 5,218,884 | 32 | 2,874,999 | 6 |
| 5 | HDPE | dockyard | *Serratia liquefaciens* | *Serratia liquefaciens* | 100.00 | 0.45 | 0.55 | 5,413,752 | 61 | 318,399 | 2 |
| 6 | HDPE | dockyard | *Bacillus licheniformis* | *Bacillus licheniformis* | 98.86 | 0.00 | 0.46 | 4,256,127 | 38 | 382,110 | 1 |
| 7 | HDPE | dockyard | *Enterobacter/Enterobacteriaceae* | *Enterobacter/Cedecea* | 98.78 | 0.60 | 0.56 | 5,014,388 | 240 | 160,006 | 3 |
| 8 | LDPE | dockyard | *Bacillus/Enterobacter* | *Bacillus* | 99.45 | 8.63 | 0.47 | 4,977,695 | 35 | 1,096,327 | 1 |
| 9 | LDPE | dockyard | *Enterobacter/Enterobacteriaceae* | *Enterobacter/Cedecea* | 98.78 | 0.60 | 0.56 | 5,014,653 | 245 | 160,006 | 3 |
| 10 | LDPE | dockyard | *Serratia* | *Serratia marcescens* | 99.86 | 0.15 | 0.59 | 5,218,582 | 30 | 2,874,999 | 6 |
| 11 | LDPE | dockyard | *Serratia fonticola* | *Serratia* | 99.94 | 0.29 | 0.54 | 5,756,538 | 113 | 185,130 | 1 |
| 12 | LDPE | falmouth wharf | *Enterococcus faecalis* | *Enterococcus faecalis* | 99.63 | 0.00 | 0.37 | 2,864,632 | 53 | 175,637 | 2 |
| 13 | PET | dockyard | *Serratia* | *Serratia marcescens* | 99.86 | 0.15 | 0.59 | 5,218,760 | 30 | 2,874,999 | 6 |
| 14 | PP | dockyard | *Serratia* | *Serratia marcescens* | 99.86 | 0.15 | 0.59 | 5,218,875 | 33 | 2,875,113 | 6 |
| 15 | PP | dockyard | *Bacillus licheniformis* | *Bacillus licheniformis* | 97.30 | 0.31 | 0.46 | 4,145,137 | 31 | 438,169 | 1 |
| 16 | PP | dockyard | *Klebsiella/Serratia/Enterobacter/Enterobacteriaceae* | *Enterobacter/Serratia* | 97.93 | 188.43 | 0.57 | 15,957,883 | 573 | 182,752 | 10 |
| 17 | PP | dockyard | *Serratia/Enterobacteriaceae* | - | 99.66 | 95.89 | 0.56 | 11,471,120 | 188 | 336,395 | 7 |
| 18 | PP | falmouth wharf | *Serratia* | *Serratia marcescens* | 99.86 | 0.15 | 0.59 | 5,219,474 | 35 | 2,875,113 | 6 |
| 19 | PVC | dockyard | *Serratia liquefaciens* | *Serratia liquefaciens* | 100.00 | 0.45 | 0.55 | 5,571,931 | 37 | 398,855 | 2 |
| 20 | PVC | falmouth wharf | *Serratia liquefaciens* | *Serratia* | 100.00 | 0.45 | 0.56 | 5,131,311 | 33 | 531,913 | 2 |

The most prevalent pathogen genus isolated was *Serratia* (Table 1), with several clones identified as *S. liquefaciens*, and *S. fonticola*, which are known human pathogens (59) that have previously been reported to occur commonly in soils in Cornwall using the *Galleria* enrichment method (60). Other species identified were *Enterococcus faecalis* which is known to infect humans (61), and *Bacillus licheniformis* which can cause opportunistic infections (e.g. (62)). Surprisingly, no virulence genes were detected in any of the assemblies, although we used a relatively stringent sequence similarity cut-off of 95%. However, a host of antimicrobial resistance genes belonging to a variety of classes were uncovered (Table 2). These include genes conferring resistance to clinically highly relevant antibiotics such as cephalosporins, aminoglycosides and trimethoprim (in *E. faecalis*) (Table 2).

**Table 2.**
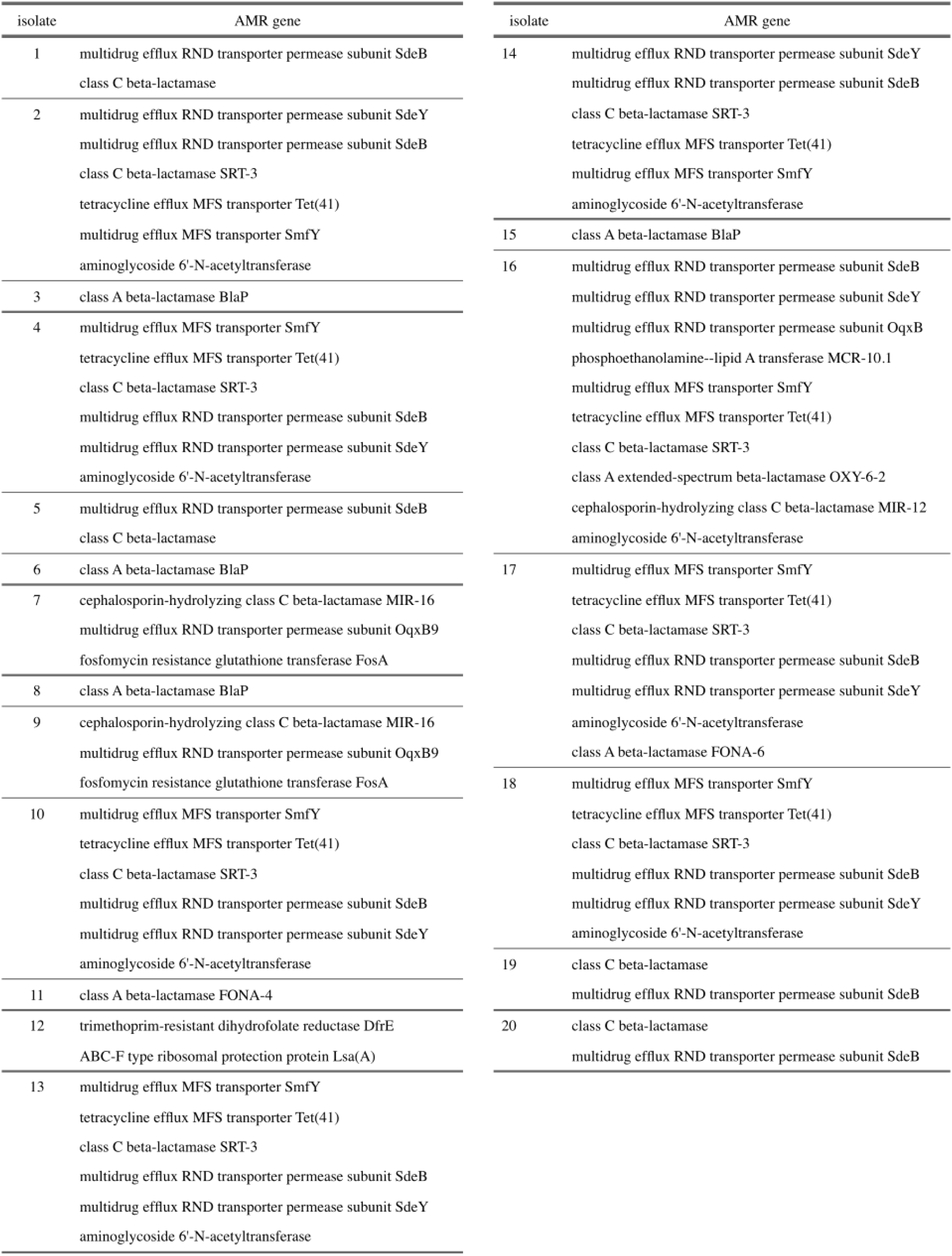
Antimicrobial resistance genes present in 20 clones isolated from plastics using the *Galleria* enrichment method.

## Discussion

There has been a recent flurry of research into the colonisation of marine plastics by bacteria, but our understanding of this process is still incomplete, especially regarding pathogenic bacteria. To help resolve ongoing debate, we submerged household plastics in the Fal Estuary (UK) and tracked bacterial colonisation as a function of plastic type, location, and time. Our results indicate that plastic colonisation in the coastal environment is largely independent of plastic type (LDPE, PE, PP, PVC and PET) with no or little effect on the number of culturable bacteria or 16S-based community composition across locations or time. The exception is LDPE, which showed lower colonisation rates in the location and colonisation experiments, although these differences were only significant compared to HDPE in the former and PET and PVC in the latter experiment (and only in the case of *E. coli* in the colonisation experiment). However, as LDPE-derived biofilms exhibited lower virulence in the *Galleria* model, it is possible that LDPE poses lower risks in terms of pathogen colonisation.

Bacterial colonisation of plastics occurred rapidly, with bacterial diversity reaching peak diversity after two weeks. Bacterial density increased with time and was highest at our final timepoint of five weeks - the only timepoint at which significant *Galleria* mortality indicated the presence of pathogenic bacteria. Our data thus suggest that total bacterial density, rather than community composition, is a better predictor of the virulence of plastic-associated biofilms. Bacterial plastic colonisation, including by pathogens, might be more rapid in summer months when higher water temperatures favour bacterial growth and persistence (63), although reduced rainfall in summer may result in lower input of terrestrial bacteria into the sea. Colonisation by bacteria likely to be of terrestrial origin (particularly those that are gut-associated and/or able to grow at 37°C on nutrient-rich medium) was more pronounced at the two locations closer to the shore and habitation, as expected. Future studies employing replicated transects could shed more light on the effect of coastal proximity on plastic colonisation.

Approaches to determine the presence of pathogenic bacteria on plastics are usually based on the ability to detect specific taxa that are selected a priori, typically via agar-based isolation (e.g. 19)). By using selective enrichment in a proven virulence model, we were able to isolate virulent bacteria, including known human pathogens such as *Enterococcus faecalis*. This species is present in marine environments (64) and has been shown to successfully colonise LDPE in lab experiments (65), but to our knowledge this is the first time that this important nosocomial pathogen has been isolated from marine plastics. The genomes recovered from plastic biofilms harboured a wide variety of antimicrobial resistance genes, highlighting the importance of plastics as a reservoir for resistance.

Plastic pollution is unabating and human exposure to micro-plastics and their associated microbial communities is likely to increase even more rapidly with a higher incidence of flooding due to climate change. Our results show that different plastic types accumulate similar bacterial communities that include a range of known human pathogens, posing a potential risk to human health.

## Acknowledgments

We are extremely grateful to Emma Walker, Timothy Bage and the boat crew from the Cornwall Port Health Authority, for kindly coordinating experimentation and data collection in the Fal Estuary, and to Macaulay Winter for assistance in the lab. Genome sequencing was provided by MicrobesNG (http://www.microbesng.com). We acknowledge support from the National Environment Research Council (NERC; grants NE/T008083/1 and NE/R011524/1). AH was supported by a Biotechnology and Biological Sciences Research Council (BBSRC) David Phillips Fellowship (BB/N020146/1).

## Supplementary Information

**Table S1.**
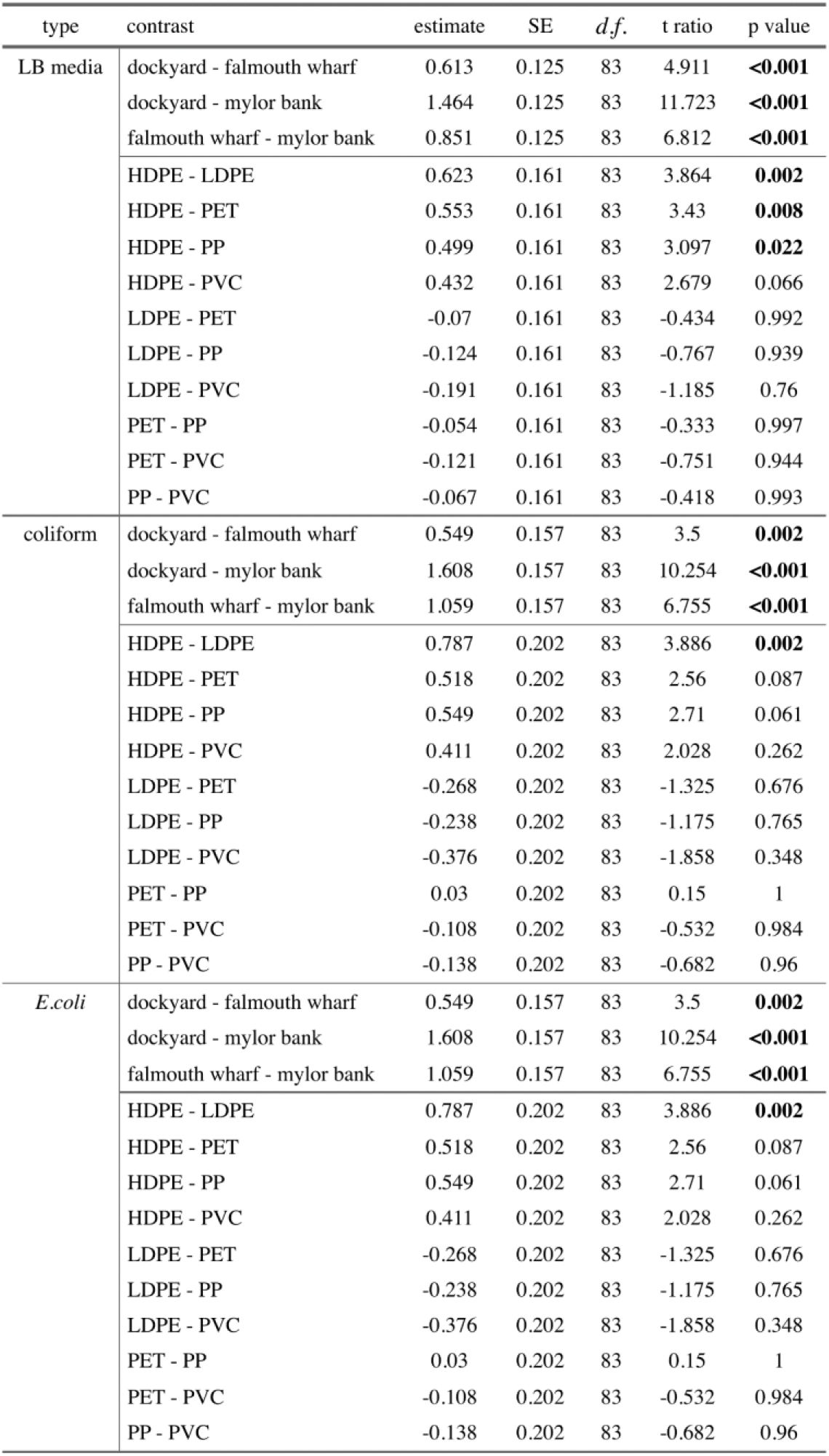
Tukey pairwise comparisons of cultural abundance of bacteria on LB media, coliform media, and *E. coli* from the location experiment. Results of multiple comparisons from the best model looking at differences between location and polymer type. Significant p values are highlighted in bold.

**Table S2.** Results of the analysis of beta-diversity of community composition between weeks and plastic type.

|  | model | *df.* | AIC | Log Lik | ANOVA<br>comparison | *F* | *P* |
| --- | --- | --- | --- | --- | --- | --- | --- |
| 1 | distance to centroid ~ week * plastic | 19 | -360.93 | 201.47 |  |  |  |
| 2 | distance to centroid ~ week + plastic | 7 | -371.21 | 194.60 | 1,2 | 1.00 | 0.451 |
| 3 | distance to centroid ~ week | 3 | -378.96 | 194.48 | 2,3 | 0.06 | 0.994 |
| 4 | distance to centroid ~ plastic | 4 | -369.41 | 190.71 | 2,4 | 2.50 | 0.063 |
| 5 | distance to centroid ~ 1 |  | -377.10 | 190.55 | 3,5 | 2.62 | 0.055 |
| 5 | distance to centroid ~ 1 |  | -377.10 | 190.55 | 4,5 | 0.07 | 0.990 |

**Table S3.** Hazard ratios between locations and plastics for both experiments. Probability above 1 is the probability that the first location/polymer type is more virulent than the baseline location/polymer type.

| contrast | experiment | Hazard ratio |  |  | Probability above 1 |
| --- | --- | --- | --- | --- | --- |
|  |  | Median | 2.5% | 97.5% |  |
| dockyard vs. falmouth wharf | location experiment | 1.00 | 0.28 | 3.76 | 0.50 |
| dockyard vs. mylor bank | location experiment | 2.57 | 0.51 | 12.98 | 0.87 |
| falmouth wharf vs. mylor bank | location experiment | 2.55 | 0.49 | 13.35 | 0.87 |
| HDPE vs. LDPE | location experiment | 4.57 | 0.88 | 23.79 | 0.97 |
|  | colonisation experiment | 1.83 | 0.46 | 6.70 | 0.80 |
| HDPE vs. PET | location experiment | 2.29 | 0.44 | 11.12 | 0.85 |
|  | colonisation experiment | 0.37 | 0.10 | 1.35 | 0.06 |
| HDPE vs. PP | location experiment | 1.99 | 0.41 | 9.04 | 0.81 |
|  | colonisation experiment | 0.87 | 0.23 | 3.27 | 0.41 |
| HDPE vs. PVC | location experiment | 1.37 | 0.19 | 9.09 | 0.63 |
|  | colonisation experiment | 0.35 | 0.10 | 1.29 | 0.06 |
| PET vs. LDPE | location experiment | 1.97 | 0.33 | 13.64 | 0.78 |
|  | colonisation experiment | 5.02 | 1.22 | 19.15 | 0.99 |
| PP vs. LDPE | location experiment | 2.31 | 0.39 | 13.73 | 0.83 |
|  | colonisation experiment | 2.05 | 0.56 | 8.27 | 0.86 |
| PP vs. PET | location experiment | 1.17 | 0.29 | 4.43 | 0.58 |
|  | colonisation experiment | 0.42 | 0.11 | 1.61 | 0.11 |
| PVC vs. LDPE | location experiment | 3.34 | 0.43 | 30.79 | 0.88 |
|  | colonisation experiment | 5.06 | 1.30 | 19.74 | 0.99 |
| PVC vs. PET | location experiment | 1.68 | 0.28 | 10.34 | 0.72 |
|  | colonisation experiment | 1.05 | 0.28 | 3.78 | 0.53 |
| PVC vs. PP | location experiment | 1.46 | 0.21 | 11.35 | 0.65 |
|  | colonisation experiment | 2.47 | 0.60 | 9.17 | 0.91 |

